# Highly efficient protein expression of *Plasmodium vivax* surface antigen, Pvs25 by silkworm, *Bombyx mori*, and its biochemical analysis

**DOI:** 10.1101/2022.03.03.482736

**Authors:** Takeshi Miyata, Kosuke Minamihata, Koichi Kurihara, Yui Kamizuru, Mari Gotanda, Momoka Obayashi, Taiki Kitagawa, Keita Sato, Momoko Kimura, Kosuke Oyama, Yuta Ikeda, Yukihiro Tamaki, Jae Man Lee, Kozue Sakao, Daisuke Hamanaka, Takahiro Kusakabe, Mayumi Tachibana, Hisham R. Ibrahim

## Abstract

*Plasmodium vivax* ookinete surface protein, Pvs25 is a transmission-blocking vaccine (TBV) candidate for malaria. Pvs25 has four EGF-like domains containing 22 cysteine residues forming 11 intramolecular disulfide bonds and this structural feature makes recombinant expression of Pvs25 difficult. In this study, we report the high expression of recombinant Pvs25 as a soluble form in silkworm, *Bombyx mori*. The Pvs25 protein was purified from hemolymphs of larvae and pupae by affinity chromatography. In the Pvs25 expressed by silkworm, no isoform with inappropriate disulfide bonds was found, requiring no further purification step which is necessary in case of *Pichia pastoris* based expressions systems. The Pvs25 from silkworm were confirmed to be the molecularly uniform by sodium dodecyl sulfate gel electrophoresis analysis and size exclusion chromatography analysis. To examine the immunogenicity, the Pvs25 from *B. mori*, was administered to BALB/c mice by the subcutaneous (s.c.) route with the oil adjuvant. The Pvs25 produced by silkworm induced potent and robust immune response, and the induced antisera correctly recognized *P. vivax* ookinetes *in vitro*, demonstrating the potency of Pvs25 from silkworm as a TBV candidate for malaria. This is the first study that to construct a mass production system for malaria TBV antigens by the silkworm to the best of our knowledge.

## Introduction

Malaria is one of the most severe arthropod-borne parasitic infectious diseases with high mortality and morbidity, especially in tropical regions of the world. The disease causes 203–262 million clinical cases per year, and the estimated annual mortality exceeds 4.0 million [1]. In particular, older adults or children are at increased risk of symptom severity. 70% of the mortality occurring in under five years children [1]. A number of malaria control measures, including chemotherapy, insecticide-treated mosquito nets (ITNs), indoor residual spraying (IRS), or paint have implemented and have contributed significantly to the reduction of malaria cases in endemic regions [1-3]. However, these control measures are suboptimal, and more the appearance of insecticide-resistant mosquitoes or antimalarial drug resistance parasite makes it difficult to control malaria [4].

*Plasmodium falciparum* causes the highest mortality rates among the five *Plasmodium* species known to infect humans. Although *Plasmodium vivax* has less mortality than *P. falciparum, P. vivax* is the most widespread human malaria and has a substantial economic impact on society [5]. In recent years, re-emergence of *P. vivax* in eradicated areas, the emergence of drug-resistant strains [6], and reports of severe disease and death have become serious public health problems. Therefore, it is considered to difficult to control and eliminate of *P. vivax* from endemic regions with current tools. Hence the research and development of new tools, including drugs, diagnostic tests, insecticides, and particularly cost-effective vaccines, should be needed for local elimination and eradicating malaria from the world [7-9].

Until now, the value of vaccine development against *P. vivax* was underestimated [10-13]. For the ultimate goal of malaria eradication, vaccine development and research against *P. vivax* cannot be neglected. Several inventive vaccine candidates have been intensively studied. For example, those targeting the asexual stages, *i*.*e*., the sporozoite, hepatic, and erythrocytic, are expected to inhibit infection and to reduce the severity. On the other hand, transmission-blocking vaccines (TBVs), which target the sexual stage at which the parasite develops spores in anopheline mosquitoes, prevents the transmission of parasites from mosquitoes to humans [14-16]. TBVs do not directly protect vaccinated individuals from infection because it induces antibodies that react with the ookinete surface proteins (OSPs) of malaria parasites in the mosquito midgut. TBVs may, therefore, be useful in not only controlling malaria but also preventing the emergence and spread of drug-resistant or the other vaccine-resistant parasites [12, 15]. TBVs can become an essential strategic vaccine when used in combination with other vaccines that target different stages of malaria parasites and serving as complementary role with each other.

Pvs25 is the most abundant OSP, might representing as much as 25% of the total OSPs. The Pvs25 function is essential for the survival of ookinetes in the mosquito midgut. Therefore, it has been studied for a promising TBV candidate against *P. vivax* [17]. The Pvs25 protein contains 4 epidermal growth factor (EGF)-like domains (EGF1 – EGF4), and glycosyl-phosphatidylinositol (GPI) anchor at the C-terminus. Despite the relatively small molecular size of Pvs25 (about 25kDa), there are 22 cysteine residues in its molecule, forming 11 disulfide bonds, making Pvs25 a complex and tangled structure [17, 18]. The recombinant expression of Pvs25 has been studied in a variety of heterologous expression systems including, *E. coli* [19], *Saccharomyces cerevisiae* [20], plant [21], and baculovirus [22]. We have also previously expressed recombinant Pvs25 using *Pichia pastoris*[23] and adenovirus[24]. In each case, the induced antiserum by recombinant Pvs25 retained the transmission-blocking (TB) activity. Since authentic Pvs25 protein has no glycosylation site, recombinant Pvs25 is not affected by post-translational modification of various expression systems. However, an advanced expression system is required to express Pvs25 with correctly connected disulfide bonds. Indeed, formation of Pvs25 isoforms due to misfolding has been observed in several expression systems [20, 23] and it is critical to yield correct structure of Pvs25 called Pvs25(A-form) that retains TBV activity. In addition, large-scale production of correctly-folded Pvs25 is essential for its development as a TBV [25].

The silkworm-baculovirus expression vector system (silkworm-BEVS) has the advantage of being suitable for the production of proteins that are challenging to produce with other protein expression systems. Particularly, the silkworm-BEVS is better at post-translational modifications than that of prokaryotes. In the cases of multimeric proteins or proteins with complex structures, such as the ones containing multiple disulfide bonds, proper folding of proteins expressed recombinantly is troublesome, and often results in formation of insoluble proteins at the bacterial expression systems. Furthermore, even if the protein is folded correctly, the expression level may not be sufficient. Silkworm BEVS has the ability to express recombinant proteins in a soluble form at relatively high expression levels [26]. For example, it has been reported that the expression level of silkworm adults is 10 to 100 times higher than that of insect cells [26]. Moreover, in the silkworm BEVS, one silkworm acts as a single cultivator, so that the production scale can be increased linearly without doing any optimization by simply increasing the number of silkworms used. The other expression systems that use liquid-culturing require optimization of culturing condition at every scaling up process to achieve high expression level and this often becomes problems. Thus, silkworm-BEVS has the potential to provide the desired recombinant protein with high productivity and at a low cost, which is suitable for production of antigen proteins for vaccines [27]. For example, we have also succeeded in the mass production of virus-like particles that form multimeric forms or surface proteins of the virus using BEVS [28, 29].

There have been several reports regarding the expression of other malaria parasite surface proteins in silkworms with retained physiological functions, however no attempt has been conducted on Pvs25 so far [30, 31]. In the present study, we report that silkworm expression system can efficiently produce the surface protein of Pvs25 from *Plasmodium vivax* and the obtained Pvs25 has excellent properties as TBV candidates in terms of homogeneity and immunogenicity.

## Materials and Methods

### Silkworm cells and strain

The NIAS-Bm-Oyanagi2 (BmO2: kindly provided from Dr. Imanishi) cells were maintained in IPL41 insect medium (Sigma, St. Louis, MO) with 10% fetal bovine serum (Gibco, Grand Island, NY) at 27°C. The d17 strains of silkworm used in this study were provided by the Institute of Genetic Resources at Kyushu University, and the silkworm larvae were reared on mulberry leaves at 24–29°C.

### Construction of silkworm-baculovirus vectors for Pvs25 proteins

To construct *Bombyx mori* expression plasmid for Pvs25, the coding region (Ala_23_–Leu_195_) was amplified by PCR from plasmid pPvs25H [23], with two primer pairs: 5′-TACTTCCAGGGAG-ATGCCGTCACGGTAGACAC -3′, and 5′-CTGGGTCTAACTCGATTAAAGGCATACATTTTTCTCT-TTGTCG -3′, to construct N-terminal tag type and 5′-GCAAGCAACGCTGATGCCGTCACGGTAG-ACAC -3′ and 5′-TTCGCCTGCACTCGAAAGGCATACATTTTTCTCTTTGTCG -3′, to construct C-terminal tag type. The amplified DNA fragment by high-fidelity DNA polymerase (New England BioLabs, Beverly, MA, USA) was purified using a PCR Purification Kit (QIAGEN Inc., Valencia, CA, USA). The fragment was subcloned into the multicloning site of the *B. mori* entry vector pENTR11L21H8STREPTEV or pENTR11L21TEVH8STREP [27] by using In-Fusion reaction to construct the entry plasmids (pBm/Pvs25-HS or pBm/HS-Pvs25) (**Fig. 1a**).

**Figure 1.**
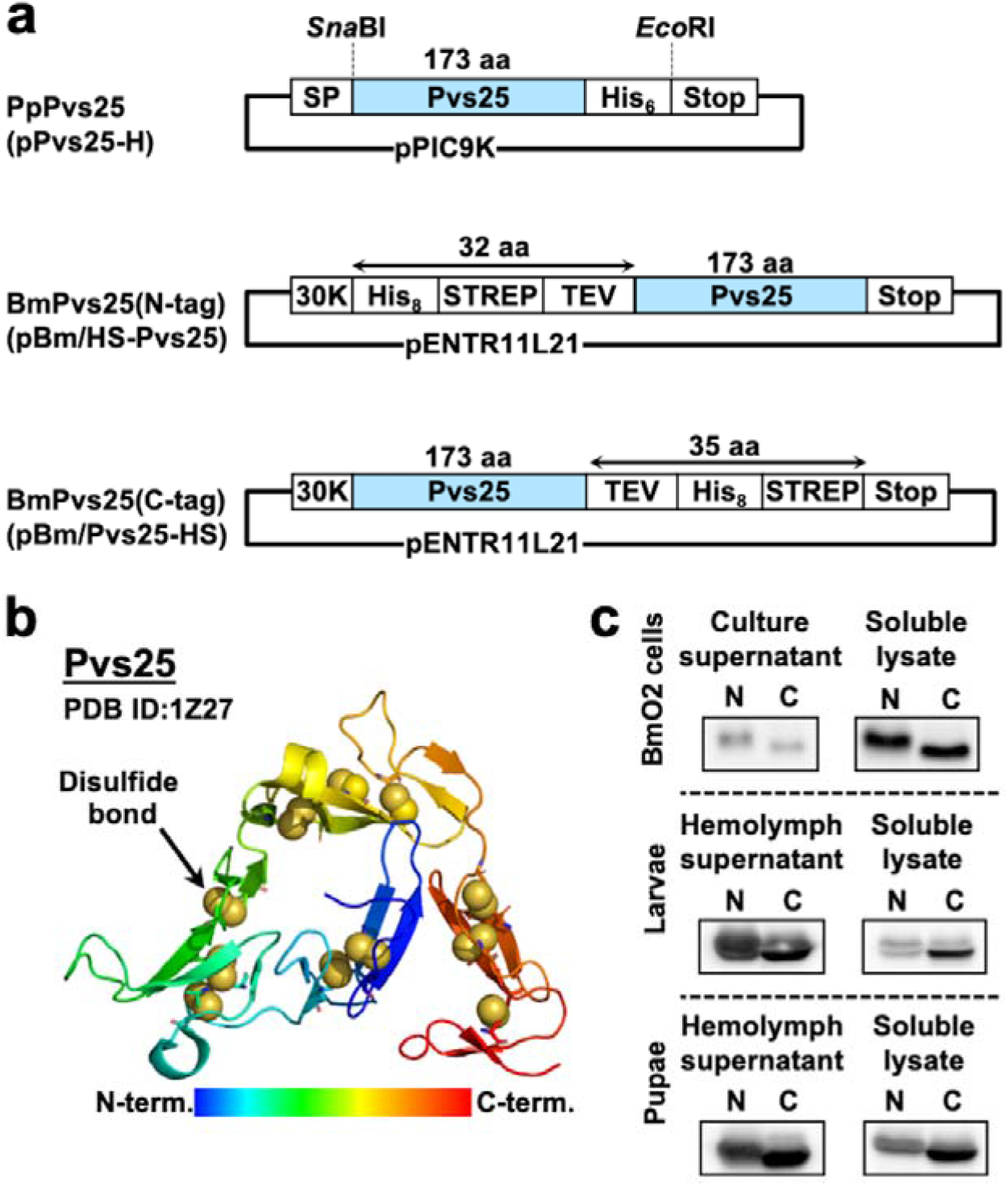
Constructs of Pvs25 used in this study and expression of the Pvs25 in *Bombyx mori*. (a) Schematic drawing of constructs of Pvs25 for *Pichia pastoris* and *B. mori* used in this study. PpPvs25 from pPvs25-H construct with C-terminal hexa-histidine-tag (His_6_) was expressed in *P. pastoris* as previously reported [23]. For the silkworm expression, The Pvs25 gene fused to octa-histidine-tag (His_8_) and Strep-Tag II (STREP: amino acid sequence of WSHPQFEK) along with tobacco etch virus protease recognition sequence (TEV: amino acid sequence of ENLYFQG) at its N or C-terminus was inserted into the multi-cloning site of pENTR11L21 entry vector by seam-less cloning method using In-Fusion [27]. (b) The 3-dimensional structure of Pvs25 showing its 11 disulfide bonds in space-filling model. The image was produced by using Pymol from the PDB data of 1Z27. (c) Western blotting analysis of expression confirmation on Pvs25 in BmO2 cells, silkworm larvae and pupae. N and C represent Pvs25 with purification tags at N-and C-terminal, respectively. The proteins were detected by targeting His_8_ using HisProbe-HRP conjugate.

pBm/HS-Pvs25 or pBm/Pvs25-HS which was designed to express Pvs25 with silkworm 30 kDa protein signal peptide (MRLTLFAFVLAVCALASNA, 30K), octahistidine-tag (His_8_), Strep-Tag II (WSHPQFEK, STREP), and tobacco etch virus (TEV) protease cleavage sequence for purification tag removal as needed (Fig. 1a). These entry plasmids were transposed to the destination vector (pDEST8, Invitrogen, USA) with Gateway LR reaction according to the manufacture’s instruction for construct transfer plasmids. The obtained transfer plasmids were transformed into *E. coli* BmDH10Bac to generate recombinant bacmid DNAs (BmNPV) [32]. The bacmid DNAs were transformed into NIAS-Bm-oyanagi2 cells with transfection reagent. The generation and amplification of recombinant baculoviruses were conducted as the protocols described previously [29]. Final viral titers were determined by the end-point dilution assay. All recombinant DNA experiments were conducted according to the Safety Guidelines for Gene Manipulation Experiments of the Kagoshima University and Kyushu University.

### Expression of recombinant Pvs25s in cultured insect cells and silkworm (BmPvs25(N-tag) and BmPvs25(C-tag))

The expression level of Pvs25s by a viral infection in NIAS-Bm-oyagami2 cells was measured as follows: 1 × 10^6^ cells per well in six-well tissue culture plates were infected with recombinant Pvs25 expressing BmNPVs at a multiplicity of infection (MOI) values of 1. At three days post-infection (dpi), after centrifugation at 1,000 x g for 10 min at 4°C, the culture supernatant (secreted proteins) was collected and cells were washed with phosphate-buffered saline (PBS) and lysed with sonication by handy mixer in 1x T buffer (20 mM Tris-HCl, 0.5 M NaCl, pH7.5) containing 0.5% CHAPS. After centrifugation at 9,600 x g for 30 min at 4°C, supernatant (intracellular proteins) was collected. The secreted proteins (5 µL) and the intracellular proteins (5 µL) were subjected to 12.5% sodium dodecyl sulfate polyacrylamide gel electrophoresis (SDS-PAGE)/western blotting analysis. After blotting to PVDF membrane, the recombinant Pvs25 proteins were detected with HisProbe-HRP conjugate by targeting His_8_ in the constructs of BmPvs25s (1:2,000 dilution, Thermo Fisher Scientific Inc., MA, USA) as previously described [27].

To investigate the expression level of Pvs25 in silkworm larvae or pupae, the recombinant baculoviruses were injected into the hemocoel of silkworm larvae (d17 strain) or pupae (d17 strain) at the dose of 1 ×10^5^ pfu per larva of pupa. After 4 dpi, the hemolymph of larvae was directly collected using a microliter syringe and centrifuged at 10,000 × g for 30 min at 4°C to remove cells in hemolymph. The larvae were then dissected to remove digestive canal and the fat body cells of larvae were collected by scraping off from the remaining skin of silkworm larvae. Pupae, on the other hand, were smashed in PBS(-), followed by centrifugation (10,000 x g for 30 min at 4 °C) to separate fat body cells which contain intracellular proteins, and the supernatant was collected as hemolymph of pupae. The remaining fat body cells of pupae were washed with PBS(-) and lysed similarly same as described above. The Pvs25 proteins from hemolymph or fat body cells of larvae and pupa were detected SDS-PAGE/western blotting analysis in the same way as described above.

### Purification of recombinant Pvs25s

Pooled hemolymph supernatant (approximately 10 mL from 20 larvae) was collected by centrifugation (9,600 × *g*) for 30 min and diluted to 50 mL with 5 x T-buffer (100 mM Tris-HCl, 2.5 M NaCl, pH7.5) containing a small amount of thiourea for inhibition of browning reaction of silkworm hemolymphs. In the case of pupae, 20 pupae were smashed with spatula and the cells were separated by centrifugation at 13,400 × *g* for 30 min to obtain the hemolymph (approx. 8 mL) and the hemolymph was diluted with 5 x T-buffer to 50 mL with adding a small amount of thiourea. The obtained samples (50 mL) were centrifuged (13,400 × *g*, 30 min), followed by filtration (0.45 µm, Minisart® Syringe Filter) to remove residual debris completely was used as a crude extract. The crude extract was applied to an immobilized metal ion affinity chromatography column (HisTrap FF, 5 mL resin; GE Healthcare, Little Chalfont, UK). After washing with 25 mL of the 1 x T-buffer containing 25 mM imidazole, the Pvs25 protein was eluted with 25 mL of the 1 x T-buffer containing 100 or 500 mM imidazole, respectively. The affinity-purified protein was then subjected to StrepTrap HP column (GE Healthcare) to remove silkworm-derived proteins and eluted with buffer containing 2.5 mM desthiobiotin (DTB) [27].

Besides, to investigate the hydrophobic profile of Pvs25 proteins, the affinity-purified Pvs25 protein was applied to hydrophobic interaction chromatography (HIC) (HiTrap Phenyl HP; 5 mL resin, GE Healthcare) In detail, the diluted Pvs25 sample (0.1 mg/mL) was adjusted to contain 2 M ammonium sulfate, bound to the column. After washing with 2 M ammonium sulfate solution (25 mL) and eluted with 25 mL of the 1 M ammonium sulfate solution in 5 mL fractions. Then, the eluate including Pvs25 protein was concentrated and replaced with PBS by ultrafiltration membrane (Amicon Ultra 15, 10 kDa MWCO).

The PpPvs25(A-form) protein was expressed in *Pichia pastoris* GS115 and purified as previously described [23].

All protein samples concentration were quantified with the BCA protein assay kit (ThremoFisher Scientific, Waltham, USA). 5,5_-dithio*bis*-(2-nitrobenzoic acid) (Ellman’s reagent; Pierce) and Protein Redox State Monitoring Kit (SulfoBiotics, Shifter, Dojindo, Japan) were used to estimate the amounts of free thiol groups.

### Size-exclusion chromatography

The purified protein was used for size-exclusion chromatography with a flow rate of 0.8 mL/min (HiLoad 16/60 Superdex 75 pg column; GE Healthcare) to evaluate the molecular weight of various recombinants Pvs25. Molecular weight standards (Gel Filtration Calibration kits LMW; GE Healthcare) were used to estimate the molecular weight of Pvs25 by calculating the partition coefficient (K_av_) values for each protein standard and sample protein. This highly purified protein sample was used for an immunization experiment.

### Immunization with recombinant Pvs25 protein and analysis of induced antibodies

Seven-week-old female BALB/c mice were purchased from Japan SLC (Fukuoka, Japan). Mice were immunized *via* the s.c. route with 30 µg of Pvs25. Incomplete Freund’s adjuvant (IFA; Difco Laboratories, Detroit, MI, USA) was used for s.c. adjuvants. The mice were immunized three times, at weeks 0, 2, and 4. Mice were anesthetized 2 weeks after the third immunization (week 4) by intraperitoneal injection of pentobarbital sodium salt (Nacalai Tesque, Inc.), and were sacrificed by exsanguination to collect serum. For specific serum antibody analysis, ELISA plates (Sumilon; Sumitomo Bakelite Co.) were coated with Pvs25 (5 µg/mL) in bicarbonate buffer (pH 9.6 by incubating the plate at 4 °C overnight. The plate was washed 3 times with PBS-T and then 3 times with PBS. The plate was blocked with 1% bovine serum albumin (BSA) in PBS for one h at 37 °C. Two-fold serial dilutions of the antisera starting with 50-fold dilution with 0.5% BSA in PBS were applied to the wells in duplicate, which were then incubated for 1 h at 37°C. Then, the plate was incubated with anti-mouse IgG conjugated to alkaline phosphatase (1/4,000; Sigma-Aldrich) for 1 h at 37 °C. *p*-Nitrophenyl phosphate solution (Bio-Rad) was added to the plate for color development, and the absorbance at OD_405_ was measured after 20 min incubation at 37 °C, using a microplate reader (Thermo Fisher Scientific, Multiskan FC). The antibody titer was defined as previously described [33].

All animal experimental protocols were approved by the Institutional Animal Care and Use Committee of the Kagoshima University, and the experiments were conducted according to the Ethical Guidelines for Animal Experiments of the Kagoshima University.

### Detection of native Pvs25 in antisera from immunized mice

Peripheral blood from *P. vivax*-infected patients was collected as described previously [3]. The gametocytemic patient blood was used to grow zygotes and ookinetes *in vitro*, as described previously [3]. They were spotted onto slides and fixed with acetone, as described previously [34-36]. The slides were blocked with 5% nonfat milk in PBS and incubated with mouse antisera derived from immunization with Pvs25 protein emulsified with IFA after diluting the antisera 100-fold with 5% nonfat milk in PBS. The samples were washed with ice-cold PBS and incubated with Alexa Fluor 488-conjugated anti-mouse antibody (Thermo Fisher Scientific). After washing with ice-cold PBS, slides were viewed by confocal scanning laser microscopy (LSM5 PASCAL; Carl Zeiss MicroImaging, Thornwood, NY, USA).

### Statistical analysis

Statistical analyses were conducted by the Kruskal-Wallis test or the Wilcoxon-Mann-Whitney test using JMP software version 8.0 (SAS Institute, Inc., Cary, NC, USA).

## Results and discussion

### Expression and purification of recombinant Pvs25 protein from *B*. *mori*

**Figure 1b** shows the crystalline structure of Pvs25 produced from the data of PDB ID of 1Z27. In the extracellular domain of Pvs25, there are 22 Cys residues (shown as space-filling model in Figure 1b) forming 11 intracellular disulfide bonds which are essential for overall structural integrity and native antigenicity of Pvs25. First, we evaluated the expression of Pvs25 by using silkworm-BEVS, which has a history of expressing proteins with complex structures [20, 37], for validation. First, we introduced the purification tags at the either N-or C-terminal of Pvs25 to examine the effect of site of purification tag on the expression level of Pvs25 in silkworm-BEVS. Both N-and C-termini are exposed on the surface of Pvs25 (Figure 1b), suggesting that the purification tags at both N-and C-termini would function. Also, in our previous report regarding expression of Pvs25 in *Pichia pastoris*, we had already demonstrated the purification of Pvs25 by using affinity-tags and the antigenicity of Pvs25 with the affinity tags remained intact [23]. The western blotting analysis results of expression of Pvs25s with N-or C-terminal purification tags in BmO2 cells and silkworm larvae and pupae were shown in **Figure 1c**. Since all constructs of Pvs25 possess 30K secretory signal peptide, the expressed Pvs25s were expected to be secreted out from cells. The Pvs25s with N-or C-terminal purification tag were successfully expressed in culture supernatant in BmO2 cells and hemolymph in silkworm larvae and pupae. Also, Pvs25s were present inside of BmO2 cells and silkworm cells as soluble form. The expression levels of Pvs25s with N-or C-terminal purification tags were comparable each other, suggesting that the position of purification tags in Pvs25 has no critical effect on the expression level of Pvs25.

Since the expression of Pvs25 in silkworm was confirmed, we proceeded to purify Pvs25 from larvae and pupae (Here after BmPvs25). We used hemolymph from 20 silkworm larvae or 20 pupae to perform purification of BmPvs25 (**Figure 2**). The detailed analysis results on the samples of purification step of Ni-affinity column chromatography were shown in **Figure S1**. In this chromatography, BmPvs25 can be eluted remarkably at 100 mM elution concentration by imidazole, but silkworm-derived proteins were also eluted at the same condition. Since it is known from previous studies that silkworm-derived proteins (about 70 kDa) bind with Ni-immobilized metal affinity resin [38], we further purified BmPvs25 with StrepTactin affinity column. The fractions of Ni-affinity chromatography containing BmPvs25 were collected (lanes 4 in Figure 2) and applied to StrepTactin column. The silkworm-derived proteins that could not be separated by Ni-affinity chromatography were successfully removed by StrepTactin column and both N-tag and C-tag types of BmPvs25s were purified efficiently (lanes 7 in Figure 2). Bands corresponding to the Pvs25s were observed in the flow-through and wash fractions as well (lanes, 2, 3, 5 and 6 in Figure 2) probably due to the overload of Pvs25 into the columns and these leaked Pvs25 could be purified by conducting chromatography again (data not shown). The purity of BmPvs25s were quite high except that BmPvs25(N-tag) expressed in pupae showed some extra bands at high molecular weight region. These contaminating proteins attached to both Ni-affinity column and StrepTactin column, implying that they possess both His_8_-tag and Strep-Tag II. Since the SDS-PAGE analysis was conducted under reducing condition, these proteins were unlikely to be the oligomers of BmPvs25(N-tag) connected via disulfide bonds. Therefore, although we didn’t further examine the features of the contaminating proteins, they may be the products of frameshifts or skipped stop codon reads during translation. Comparing the purification efficiency between larvae and pupae, although the hemolymph of larvae contained more silkworm-derived proteins than hemolymph of pupae, the overall purity of BmPvs25s were comparable each other, resulting a single band after 2 steps of affinity chromatography, except BmPvs25(N-tag) expressed in pupae mentioned above. The overall yields of BmPvs25s were 0.6 ± 0.1 mg/larva or 1.2± 0.4 mg/pupa and there were no significant differences between N-tag and C-tag versions of BmPvs25.

**Figure 2.**
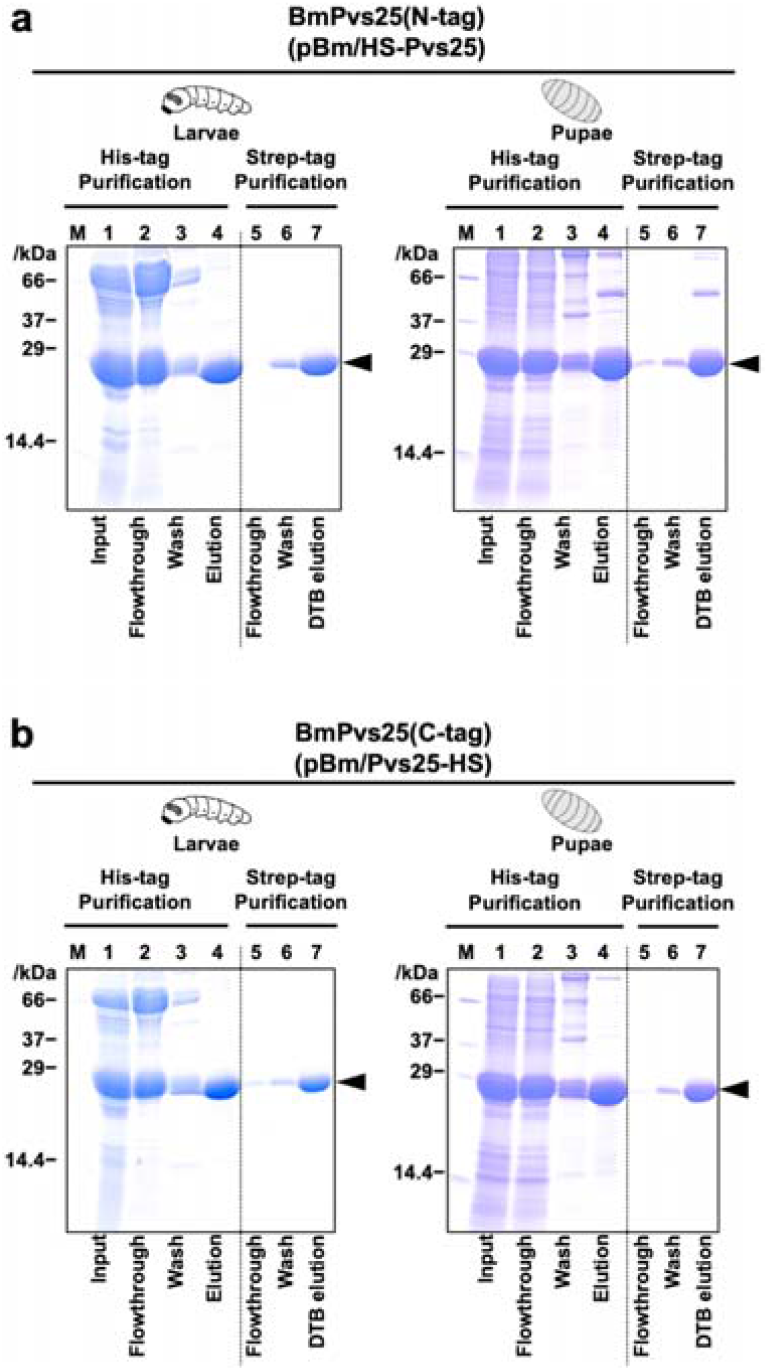
Purifications of BmPvs25s expressed in silkworm larvae and pupae using Ni-affinity chromatography and Strep-Tag II purification. (a) Purification results of BmPvs25(N-tag). (b) Purification results of BmPvs25(C-tag). Purified BmPvs25s were analyzed by SDS-PAGE (12.5% acrylamide)/CBB-staining. M, molecular marker; lane 1, intact protein from hemolymph of larvae or pupae; lane 2, flowthrough fraction of Ni-affinity column (HisTrap excel); lane 3, wash fraction of Ni-affinity column washed with 1 x T-buffer containing 25 mM imidazole; lanes 4, Ni-affinity chromatography elution fraction (eluted with) after dialysis and concentration with ultrafiltration membrane; lane 5 flowthrough fraction of StrepTrap HP column; lane 6, wash fraction of StrepTrap HP column washed with 1 x T-buffer; lane 7. Elution fractions eluted with 1 x T-buffer containing 2.5 mM desthiobiotin (DTB). The positions of the BmPvs25s protein were shown by arrowhead on the right, respectively.

Because of the complex structure of Pvs25, *E. coli* expression systems was not able to express Pvs25 as soluble protein, resulting in formation of inclusion body [19, 37, 39]. Therefore, yeast *Saccharomyces cerevisiae* expression systems [18, 20, 39, 40] or *Pichia pastoris* [23] has been used. In the yeast expression system, Pvs25 can be expressed as a soluble protein, however, the isoforms of Pvs25 were produced due to a misconnection of disulfide bonds, which does not function as a vaccine antigen [20]. The expression in *P. pastoris*, which we had previously validated, was more productive and useful than in baker’s yeast, but the proportion of isoform remained high as before, requiring additional purification steps to separate correctly folded isoform, hereafter termed as Pvs25(A-form). In this context, we subsequently analyzed the contents of isoforms in BmPvs25s.

### Isoform analysis on BmPvs25s

The structure of the correctly folded Pvs25(A-form) is rigid by being cross-linked with 11 disulfide bonds and holds the hydrophobic amino acid residues in place. Due to this feature of Pvs25(A-form), it is stable against heat treatment under non-reducing condition. However, once the disulfide bonds in Pvs25 are reduced, the structure of Pvs25 will be disrupted even without heat treatment and the intrinsic hydrophobic residues will be exposed, resulting in a great increase of the mobility of bands in SDS-PAGE analysis [23]. While incorrectly folded Pvs25 (termed as B-form) [23] exposes its hydrophobic residues without doing heat or reducing treatment, thus the mobility of its band is higher than Pvs25(A-form). Therefore, we evaluated the mobility of bands of BmPvs25s in SDS-PAGE after heat and/or reducing treatment to assess the content of A-and B-forms of Pvs25. (**Figure 3**).

**Figure 3.**
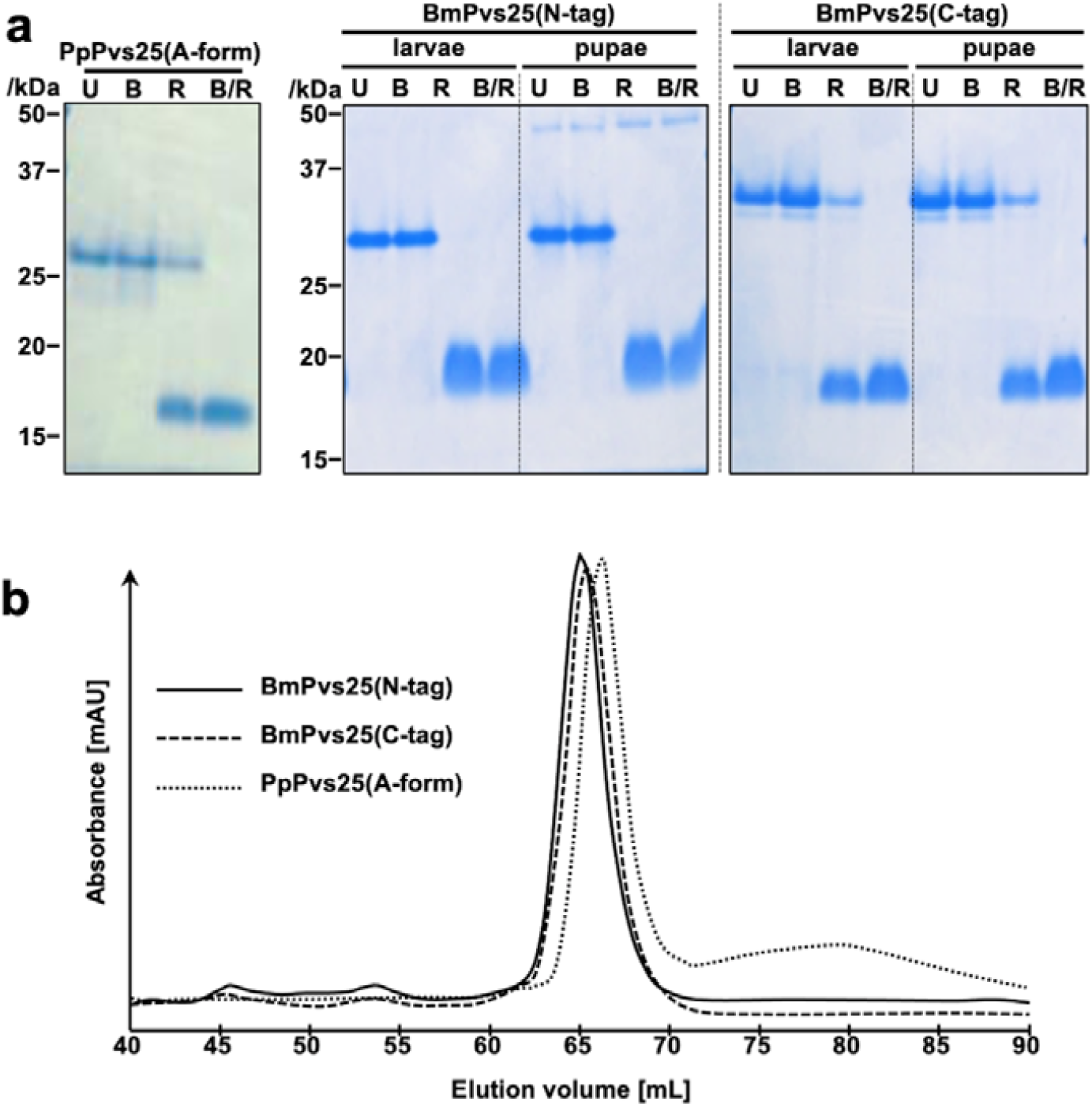
Isoform analysis of BmPvs25s. (a) The affinity-purified Pvs25s (PpPvs25(A-form), BmPvs25(N-tag) or BmPvs25(C-tag)) were subjected to 12.5% acrylamide SDS-PAGE under various conditions. U: untreated, B: boiled under non-reducing condition, R: reducing condition by addition of DTT without heat treatment, B/R: boiled under reducing condition by addition of DTT. (b) size-exclusion chromatography analysis of BmPvs25s from silkworm larvae and PpPvs25 (A-form) under the native state.

As a comparison, PpPvs25(A-form) purified from *P. pastoris* according to the previous report was used [23]. All BmPvs25s expressed in both silkworm larvae and pupae showed the same band pattern as the correctly folded PpPvs25(A-form), indicating that BmPvs25s were composed of only the correctly folded A-form. We also checked the BmPvs25s with Ellman’s reagent to confirm the absence of free thiols (data not shown). Moreover, we conducted hydrophobic interaction chromatography analysis on the BmPvs25(N-tag) from larvae, by which the PpPvs25(A-form) can be separated from incorrectly folded isomers, and we found that BmPvs25(N-tag) were eluted at the fractions attributed to correctly folded A-form (**Figure S2**). Comparing N-tag and C-tag versions of BmPvs25s, the BmPvs25(C-tag) in lanes U and B in **Figure 3a** showed lower band mobility. This can be attributed to the molecular weight difference between BmPvs25(C-tag) (22.9 kDa) and BmPvs25(N-tag) (22.7 kDa)(See supporting information for the whole amino acid sequences of BmPvs25s). Also, Pvs25 possesses its N-terminus at the center of its planer triangle structure (**Figure 1b**), meanwhile the C-terminus is located at an apex of the triangle structure. Hence, BmPvs25(C-tag) has more larger 3-dimentional structure than BmPvs25(N-tag), resulting in lower mobility in SDS-PAGE analysis. Considering that there were no structural differences present between the BmPvs25s expressed in silkworm larvae and pupae and in the case of BmPvs25(N-tag) expressed in pupae contained contaminating proteins, we conducted following experiments with only the ones expressed with larvae. To verify the homogeneity of the BmPvs25s, size-exclusion chromatography was used (**Figure 3b**). All samples were eluted as a single peak with slightly different elution volumes, which is attributed to the molecular weight differences of PpPvs25(A-form) and BmPvs25s. From these SEC chromatograms, we conclude that BmPvs25s were homogeneous and no oligomers or aggregates of Pvs25 were present in the purified samples. We also checked whether the BmPvs25s were glycosylated or not but there were no N-type or O-type glycans attached to BmPvs25s (data not shown). This is consistent with the absence of N-type glycan recognition sequences in the amino acid sequence of Pvs25 and the absence of glycan chains in PpPvs25 [23, 40]. These results indicate that silkworm-BEVS is a suitable expression system for producing high quality Pvs25 antigen, which is a promising TBV.

### Immunogenicity assessment of BmPvs25

The immunogenicity of BmPvs25s and PpPvs25(A-form) was tested by subcutaneous administration to mice. Since IFA was used as an adjuvant for all samples, an IFA-treated group was established as a comparison group. The measurement results of IgG titers against Pvs25 in mouse sera are shown in **Figure 4**. The mice immunized with BmPvs25s produced anti-Pvs25 IgGs in the serum at comparable titer levels as the ones immunized with PpPvs25(A-form). There was no significant difference in titers between both N-tag or C-tag types of BmPvs25s and PpPvs25(A-form). In the previous report, mice immunized with PpPvs25(A-form) successfully generated anti-Pvs25 IgG that can function as TBV, therefore the anti-Pvs25 IgG induced by BmPvs25s can be also capable of working as TBV.

**Figure 4.**
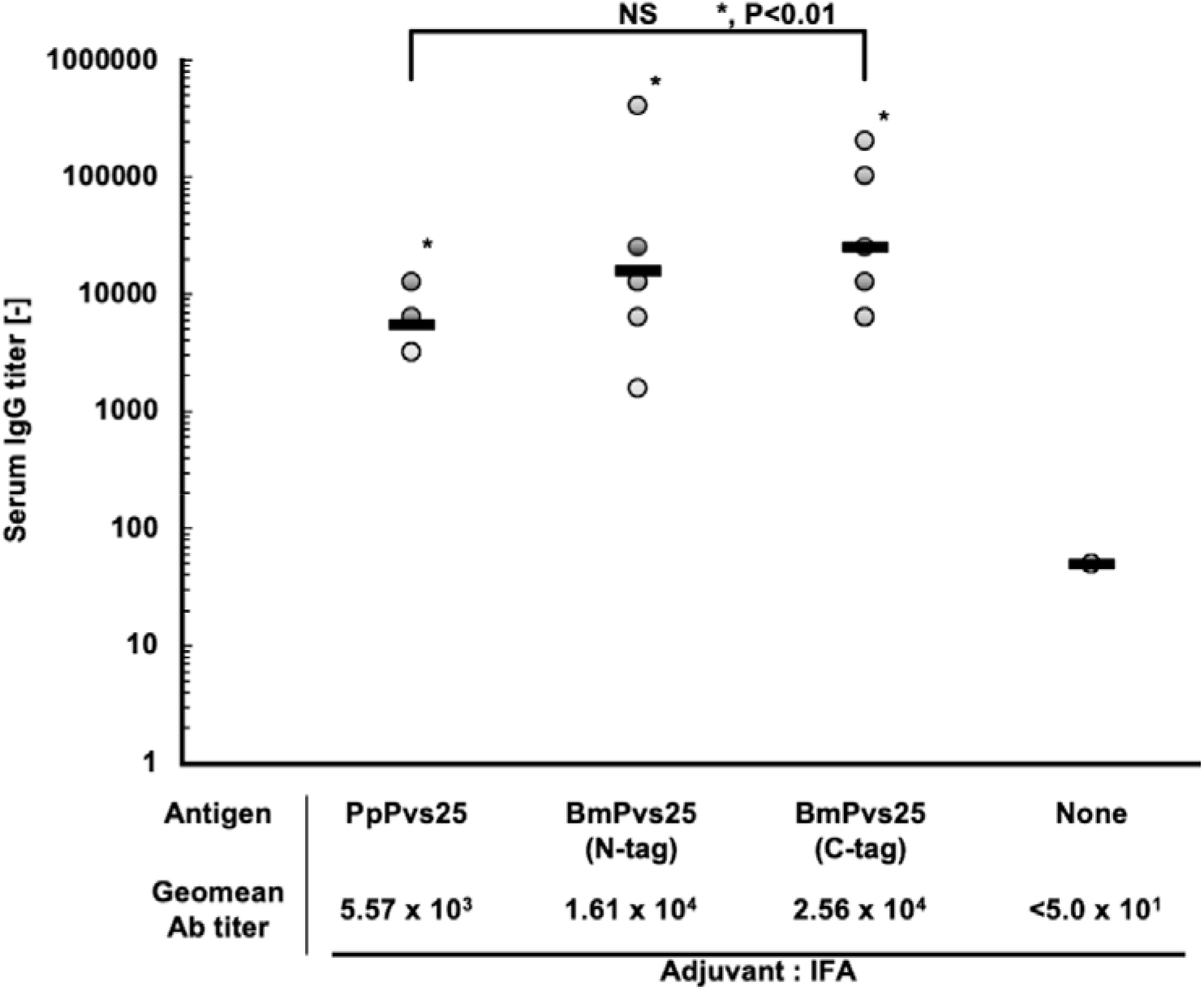
Immunogenicity of the Pvs25 proteins for induction of a Pvs25-specific serum IgG response. Female BALB/c mice (five or six mice per group) were immunized with PpPvs25 (A-form), BmPvs25(N-tag) or BmPvs25(C-tag), at 30 µg/shot of Pvs25, by the subcutaneous (s.c.), three times, at week 0, 2 and 4. Antiserum samples were collected 2 weeks after the third immunization (week 6) and were evaluated for Pvs25-specific IgG titers. Incomplete Freund’s adjuvant (IFA) was used as s.c. vaccine adjuvant. IFA alone immunized serum (None) was used as a negative control. Antibody titers were defined as the serum dilution at which OD_405_ is 0.1, or the serum dilution at which OD_405_ is < 0.1 at one higher dilution (two-fold) as described previously [23, 24, 33]. *, significantly different from negative control group by Wilcoxon-Mann-Whitney test (*P* < 0.05). NS indicates not significant among the various Pvs25 immunized groups. PpPvs25(A-form) was used as the antigen to coat the ELISA plate.

Lastly, we evaluated whether the anti-Pvs25 IgG induced by BmPvs25 in mice can function as a transmission-blocking antibody or not by conducting immunofluorescence staining of *P. vivax* (**Figure 5** and **Figure S3** for additional images). The mouse sera immunized with BmPvs25 were able to recognize *P. vivax* in the same manner as the sera immunized with PpPvs25(A-form). As a positive control, anti-Pvs25 IgG rabbit was also used. The fluorescent images stained with mouse sera induced by BmPvs25s and the RMAL154 were totally overlapped each other, indicating that anti-Pvs25 IgG in the mouse sera recognized Pvs25 on the surface of *P. vivax*. From these results, we concluded that Pvs25 prepared by silkworm-BEVS was immunologically active and could induce antibodies capable of recognizing *P. vivax*, demonstrating their potential as a TBV.

**Figure 5.**
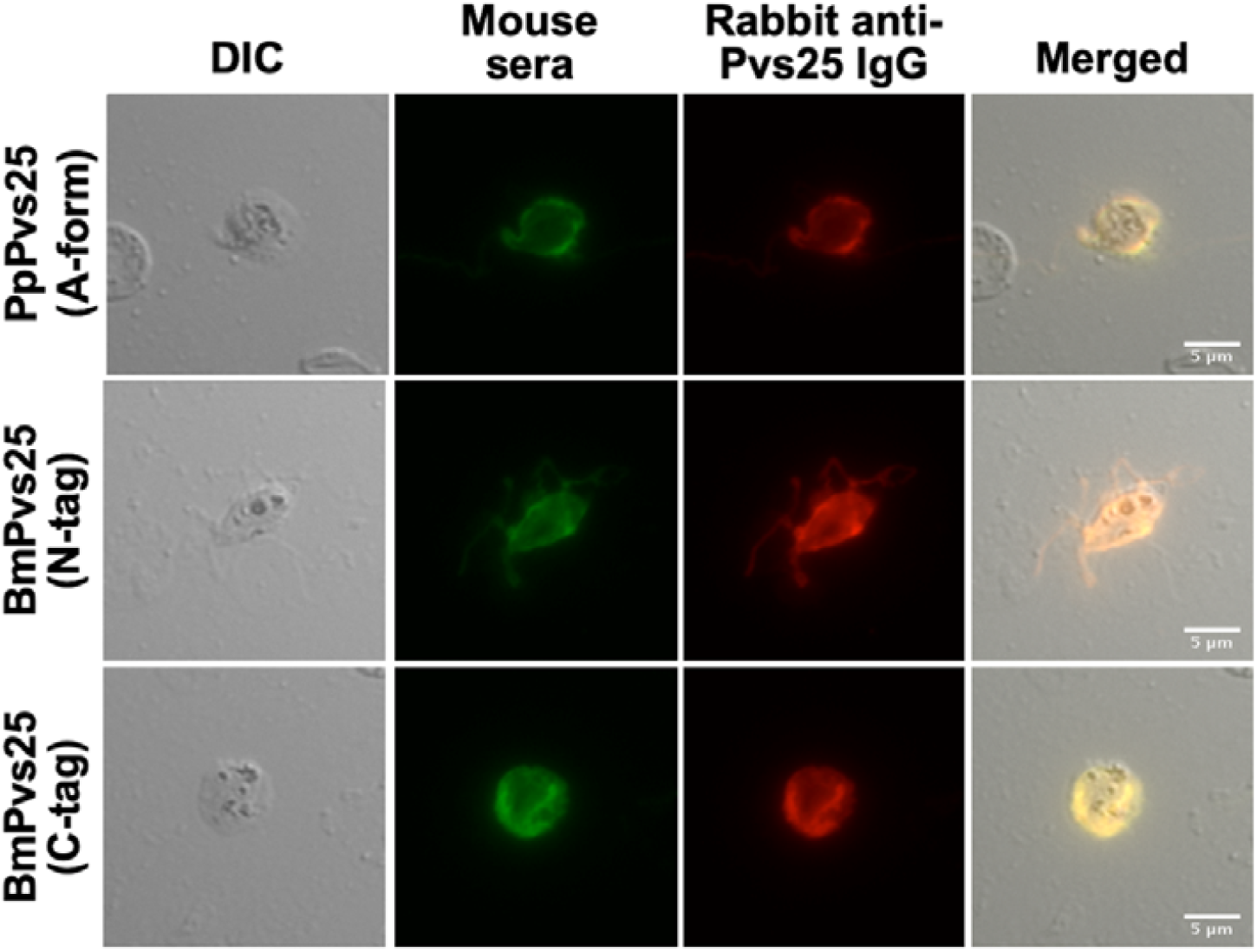
Parasite recognition of mouse sera immunized with BmPvs25s. The malaria parasite-specific reactivities of the antisera were analyzed by immunofluorescence assay. Zygotes and ookinetes (retorts and mature ookinetes) cultured *in vitro* were fixed with acetone. The immunostaining results by using mouse sera induced by PpPvs25(A-form), BmPvs25(N-tag), or BmPvs25(C-tag) were indicated as green color (Alexa Fluor 488). Immunostaining results using rabbit antibody specific to Pvs25 were indicated as red color (Alexa Fluor 546). Merged, merged image of green and red signal. Bars, 5 µm.

## Conclusion

In this study, we demonstrated for the first time the production of Pvs25, one of the major ookinete surface proteins of *P. vivax* and potential transmission-blocking vaccine, by using silkworm-BEVS. The Pvs25s with purification tags at N-or C-termini were constructed and both were efficiently expressed as soluble forms in silkworm larvae and pupae. After 2 steps of purifications using Ni-affinity column and StrepTactin column, the BmPvs25s were purified in high purity with high yields. Notably, the Pvs25s expressed in silkworm were structurally homogeneous, yielding only the correctly folded A-form isoform, which is immunologically active to induce antibodies that can block transmission of malaria. Lastly, the immunogenicity of the BmPvs25 was evaluated by immunizing mice and analyzing the antibodies in the sera. The mice immunized with BmPvs25 generated anti-Pvs25 IgG at a comparable level to the ones immunized with PpPvs25(A-form) produced by *P. pastoris*. Moreover, immunofluorescence staining of *P vivax* using mice sera immunized with BmPvs25 successfully visualized the surface antigen of Pvs25, indicating that the antibodies induced by BmPvs25 can function to block the transmission of *P. vivax*. We concluded that the silkworm-BEVS is a highly potent to produce Pvs25 for developing it as TBV for malaria.

## Acknowledgments

This work was supported by JSPS KAKENHI Grant Numbers JP17K08810, JP26460511, JP24790404.

## Notes

### Competing Interest Statement

The authors have declared no competing interest.

## References

[1] WHO, WORLD MALARIA REPORT 2018, 2018.

[2] B. Mosqueira, J. Chabi, F. Chandre, M. Akogbeto, J.M. Hougard, P. Carnevale, S. Mas-Coma, Efficacy of an insecticide paint against malaria vectors and nuisance in West Africa--part 2: field evaluation. Malar J 9 (2010) 341.

[3] N. Suwanabun, J. Sattabongkot, T. Tsuboi, M. Torii, N. Maneechai, N. Rachapaew, N. Yimamnuaychok, V. Punkitchar, R.E. Coleman, Development of a method for the in vitro production of Plasmodium vivax ookinetes. J Parasitol 87 (2001) 928–930.

[4] C. Roucher, C. Rogier, F. Dieye-Ba, C. Sokhna, A. Tall, J.F. Trape, Changing malaria epidemiology and diagnostic criteria for Plasmodium falciparum clinical malaria. PLoS ONE 7 (2012) e46188.

[5] D.K. Kochar, A. Das, S.K. Kochar, V. Saxena, P. Sirohi, S. Garg, A. Kochar, M.P. Khatri, V. Gupta, Severe Plasmodium vivax malaria: a report on serial cases from Bikaner in northwestern India. Am J Trop Med Hyg 80 (2009) 194–198.

[6] J.K. Baird, Resistance to therapies for infection by Plasmodium vivax. Clin Microbiol Rev 22 (2009) 508–534.

[7] B.M. Greenwood, D.A. Fidock, D.E. Kyle, S.H. Kappe, P.L. Alonso, F.H. Collins, P.E. Duffy, Malaria: progress, perils, and prospects for eradication. J Clin Invest 118 (2008) 1266–1276.

[8] G.A. Targett, B.M. Greenwood, Malaria vaccines and their potential role in the elimination of malaria. Malar J 7 Suppl 1 (2008) S10.

[9] B. Genton, Malaria vaccines: a toy for travelers or a tool for eradication? Expert Rev Vaccines 7 (2008) 597–611.

[10] BillandMelindaFoundation, GLOBAL HEALTH PROGRAM, 2009.

[11] MalariaVaccineTechonologyRoadmap, MalariaVaccine Techonology Roadmap, 2006.

[12] A.J. Birkett, PATH Malaria Vaccine Initiative (MVI): Perspectives on the status of malaria vaccine development. Hum vaccin 6 (2010) 139–145.

[13] M. Arevalo-Herrera, C. Chitnis, S. Herrera, Current status of Plasmodium vivax vaccine. Hum vaccin 6 (2010).

[14] T. Tsuboi, M. Tachibana, O. Kaneko, M. Torii, Transmission-blocking vaccine of vivax malaria. Parasitol Int 52 (2003) 1–11.

[15] D.C. Kaslow, Transmission-blocking vaccines: uses and current status of development. Int J Parasitol 27 (1997) 183–189.

[16] R. Carter, Transmission blocking malaria vaccines. Vaccine 19 (2001) 2309–2314.

[17] D.C. Kaslow, I.A. Quakyi, C. Syin, M.G. Raum, D.B. Keister, J.E. Coligan, T.F. McCutchan, L.H. Miller, A vaccine candidate from the sexual stage of human malaria that contains EGF-like domains. Nature 333 (1988) 74–76.

[18] A.K. Saxena, K. Singh, H.P. Su, M.M. Klein, A.W. Stowers, A.J. Saul, C.A. Long, D.N. Garboczi, The essential mosquito-stage P25 and P28 proteins from Plasmodium form tile-like triangular prisms. Nat Struct Mol Biol 13 (2006) 90–91.

[19] S.U. Moon, H.H. Kim, T.S. Kim, K.M. Choi, C.M. Oh, Y.J. Ahn, S.K. Hwang, Y. Sohn, E.H. Shin, H. Kim, H.W. Lee, Blocking effect of a monoclonal antibody against recombinant Pvs25 on sporozoite development in Anopheles sinensis. Clin Vaccine Immunol 17 (2010) 1183–1187.

[20] A.P. Miles, Y. Zhang, A. Saul, A.W. Stowers, Large-scale purification and characterization of malaria vaccine candidate antigen Pvs25H for use in clinical trials. Protein Expr Purif 25 (2002) 87–96.

[21] A.M. Blagborough, K. Musiychuk, H. Bi, R.M. Jones, J.A. Chichester, S. Streatfield, K.A. Sala, S.E. Zakutansky, L.M. Upton, R.E. Sinden, I. Brian, S. Biswas, J. Sattabonkot, V. Yusibov, Transmission blocking potency and immunogenicity of a plant-produced Pvs25-based subunit vaccine against Plasmodium vivax. Vaccine 34 (2016) 3252–3259.

[22] A.M. Blagborough, S. Yoshida, J. Sattabongkot, T. Tsuboi, R.E. Sinden, Intranasal and intramuscular immunization with Baculovirus Dual Expression System-based Pvs25 vaccine substantially blocks Plasmodium vivax transmission. Vaccine 28 (2010) 6014–6020.

[23] T. Miyata, T. Harakuni, T. Tsuboi, J. Sattabongkot, H. Kohama, M. Tachibana, G. Matsuzaki, M. Torii, T. Arakawa, Plasmodium vivax ookinete surface protein Pvs25 linked to cholera toxin B subunit induces potent transmission-blocking immunity by intranasal as well as subcutaneous immunization. Infect Immun 78 (2010) 3773–3782.

[24] T. Miyata, T. Harakuni, H. Sugawa, J. Sattabongkot, A. Kato, M. Tachibana, M. Torii, T. Tsuboi, T. Arakawa, Adenovirus-vectored Plasmodium vivax ookinete surface protein, Pvs25, as a potential transmission-blocking vaccine. Vaccine 29 (2011) 2720–2726.

[25] T. Tsuboi, S. Takeo, H. Iriko, L. Jin, M. Tsuchimochi, S. Matsuda, E.T. Han, H. Otsuki, O. Kaneko, J. Sattabongkot, R. Udomsangpetch, T. Sawasaki, M. Torii, Y. Endo, Wheat germ cell-free system-based production of malaria proteins for discovery of novel vaccine candidates. Infect Immun 76 (2008) 1702–1708.

[26] A. Usami, S. Ishiyama, C. Enomoto, H. Okazaki, K. Higuchi, M. Ikeda, T. Yamamoto, M. Sugai, Y. Ishikawa, Y. Hosaka, T. Koyama, Y. Tobita, S. Ebihara, T. Mochizuki, Y. Asano, H. Nagaya, Comparison of recombinant protein expression in a baculovirus system in insect cells (Sf9) and silkworm. J Biochem 149 (2011) 219–227.

[27] T. Mitsudome, J. Xu, Y. Nagata, A. Masuda, K. Iiyama, D. Morokuma, Z. Li, H. Mon, J.M. Lee, T. Kusakabe, Expression, purification, and characterization of endo-beta-Nacetylglucosaminidase H using baculovirus-mediated silkworm protein expression system. Appl Biochem Biotechnol 172 (2014) 3978–3988.

[28] A. Masuda, J.M. Lee, T. Miyata, T. Ebihara, K. Kakino, M. Hino, R. Fujita, H. Mon, T. Kusakabe, Stable trimer formation of spike protein from porcine epidemic diarrhea virus improves the efficiency of secretory production in silkworms and induces neutralizing antibodies in mice. Vet Res 52 (2021) 102.

[29] A. Masuda, J.M. Lee, T. Miyata, T. Sato, S. Hayashi, M. Hino, D. Morokuma, N. Karasaki, H. Mon, T. Kusakabe, Purification and characterization of immunogenic recombinant virus-like particles of porcine circovirus type 2 expressed in silkworm pupae. J Gen Virol 99 (2018) 917–926.

[30] A.L. Pang, C.N. Hashimoto, L.Q. Tam, Z.Q. Meng, G.S. Hui, W.K. Ho, In vivo expression and immunological studies of the 42-kilodalton carboxyl-terminal processing fragment of Plasmodium falciparum merozoite surface protein 1 in the baculovirus-silkworm system. Infect Immun 70 (2002) 2772–2779.

[31] V.K. Deo, Y. Inagaki, E.H. Murhandarwati, W. Asmara, T. Miyazaki, T. Kato, E.Y. Park, Sero-diagnostic potential of Plasmodium falciparum recombinant merozoite surface protein (MSP)-3 expressed in silkworm. Parasitol Int 72 (2019) 101938.

[32] T.N. Chikako Ono, Yasuyuki Nishijima, Shin-ichiro Asano, Ken Sahara and Hisanori Bando, Construction of the BmNPV T3 bacmid system and its application to the functional analysis ofBmNPV he65. Journal of Insect Biotechnology and Sericology 76 (2007) 161–167.

[33] T. Miyata, T. Harakuni, T. Tsuboi, J. Sattabongkot, A. Ikehara, M. Tachibana, M. Torii, G. Matsuzaki, T. Arakawa, Tricomponent immunopotentiating system as a novel molecular design strategy for malaria vaccine development. Infect Immun 79 (2011) 4260–4275.

[34] T. Arakawa, T. Tsuboi, A. Kishimoto, J. Sattabongkot, N. Suwanabun, T. Rungruang, Y. Matsumoto, N. Tsuji, H. Hisaeda, A. Stowers, I. Shimabukuro, Y. Sato, M. Torii, Serum antibodies induced by intranasal immunization of mice with Plasmodium vivax Pvs25 co-administered with cholera toxin completely block parasite transmission to mosquitoes. Vaccine 21 (2003) 3143–3148.

[35] T. Arakawa, A. Komesu, H. Otsuki, J. Sattabongkot, R. Udomsangpetch, Y. Matsumoto, N. Tsuji, Y. Wu, M. Torii, T. Tsuboi, Nasal immunization with a malaria transmission-blocking vaccine candidate, Pfs25, induces complete protective immunity in mice against field isolates of Plasmodium falciparum. Infect Immun 73 (2005) 7375–7380.

[36] T. Arakawa, M. Tachibana, T. Miyata, T. Harakuni, H. Kohama, Y. Matsumoto, N. Tsuji, H. Hisaeda, A. Stowers, M. Torii, T. Tsuboi, Malaria ookinete surface protein-based vaccination via the intranasal route completely blocks parasite transmission in both passive and active vaccination regimens in a rodent model of malaria infection. Infect Immun 77 (2009) 5496–5500.

[37] J.N. McCaffery, J.A. Fonseca, B. Singh, M. Cabrera-Mora, C. Bohannon, J. Jacob, M. Arevalo-Herrera, A. Moreno, A Multi-Stage Plasmodium vivax Malaria Vaccine Candidate Able to Induce Long-Lived Antibody Responses Against Blood Stage Parasites and Robust Transmission-Blocking Activity. Front Cell Infect Microbiol 9 (2019) 135.

[38] A. Morio, J. Xu, A. Masuda, Y. Kinoshita, M. Hino, D. Morokuma, H.M. Goda, N. Okino, M. Ito, H. Mon, R. Fujita, T. Kusakabe, J.M. Lee, Expression, purification, and characterization of highly active endo-α-N-acetylgalactosaminidases expressed by silkworm-baculovirus expression system. Journal of Asia-Pacific Entomology 22 (2019) 404–408.

[39] D.C. Kaslow, I.C. Bathurst, T. Lensen, T. Ponnudurai, P.J. Barr, D.B. Keister, Saccharomyces cerevisiae recombinant Pfs25 adsorbed to alum elicits antibodies that block transmission of Plasmodium falciparum. Infect Immun 62 (1994) 5576–5580.

[40] H. Hisaeda, A.W. Stowers, T. Tsuboi, W.E. Collins, J.S. Sattabongkot, N. Suwanabun, M. Torii, D.C. Kaslow, Antibodies to malaria vaccine candidates Pvs25 and Pvs28 completely block the ability of Plasmodium vivax to infect mosquitoes. Infect Immun 68 (2000) 6618–6623.

