## Supporting information for "Highly efficient protein expression of *Plasmodium vivax* surface antigen, Pvs25 by silkworm, *Bombyx mori*, and its biochemical analysis"

Running title: Bombyx mori expressed Pvs25 vaccine antigen

Takeshi Miyata[1*], Kosuke Minamihata[2], Koichi Kurihara[1], Yui Kamizuru[1], Mari Gotanda[1], Momoka Obayashi[1], Taiki Kitagawa[1], Keita Sato[1], Momoko Kimura[1], Kosuke Oyama[1], Yuta Ikeda[1], Yukihiro Tamaki[3], Jae Man Lee[4], Kozue Sakao[1], Daisuke Hamanaka[1], Takahiro Kusakabe[5], Mayumi Tachibana[6] and Hisham R. Ibrahim[1]

[1] The United Graduate School of Agricultural Sciences, Kagoshima University, 1-21-24 Korimoto, Kagoshima 890-0065, Japan; [2] Department of Applied Chemistry, Graduate School of Engineering, Kyushu University, 744 Motooka, Nishi-ku, Fukuoka 819-0395, Japan;[3] Immunobiology, Department of Infectious Diseases, COMB, Tropical Biosphere Research Center, University of the Ryukyus, 1 Senbaru, Nishihara, Okinawa 903-0213, Japan; [4]Laboratory of Creative Science for Insect Industries, Kyushu University Graduate School of Bioresource and Bioenvironmental Sciences, Motooka 744, Nishi-ku, Fukuoka 819-0395, Japan. ; [5]Laboratory of Insect Genome Science, Kyushu University Graduate School of Bioresource and Bioenvironmental Sciences, 744 Motooka, Nishi-ku, Fukuoka 819-0395, Japan. ; [6]Department of Molecular Parasitology, Ehime University Graduate School of Medicine, Shitsukawa, Toon, Ehime 791-0295, Japan.

[*]Corresponding author: Takeshi Miyata, Ph.D.
Division of Molecular Functions of Food, Department of Biochemistry and Biotechnology, Kagoshima University, 1-21-24 Korimoto, Kagoshima 890-0065, Japan

### Amino acid sequences

**PpPvs25**

MRFPSIFTAVLPAASSALAAPVNTTTEDETAQIPAEAVIGYSDLEGDFDVAVLPFSNSTNNGLLFINTTIASIAAK
EEGVSLEKREAEAAVTVDTICKNGQLVQMSNHFKCMCNEGLVHLSENTCEEKNECKKETLGKACGEFGQCIENPDP
AQVNMYKCGCIQGYTLKEDTCVLDVCQYKNCGESGECIVEYLSETKSAGCSCAIGKVPNPEDEKKCTKTGETACQL
KCNTDNEVCKNVEGVYKCQCMEGFTFDKEKNVCLGPGPHHHHHH

**BmPvs25(N-tag)**

MRLTLFAFVLAVCALASNAHHHHHHHHGGGSAWSHPQFEKGGGENLYFQGDAVTVDTICKNGQLVQMSNHFKCMCN
EGLVHLSENTCEEKNECKKETLGKACGEFGQCIENPDPAQVNMYKCGCIEGYTLKEDTCVLDVCQYKNCGESGECI
VEYLSEIQSAGCSCAIGKVPNPEDEKKCTKTGETACQLKCNTDNEVCKNVEGVYKCQCMEGFTFDKEKNVCL

**BmPvs25(C-tag)**

MRLTLFAFVLAVCALASNADAVTVDTICKNGQLVQMSNHFKCMCNEGLVHLSENTCEEKNECKKETLGKACGEFGQ
CIENPDPAQVNMYKCGCIEGYTLKEDTCVLDVCQYKNCGESGECIVEYLSEIQSAGCSCAIGKVPNPEDEKKCTKT
GETACQLKCNTDNEVCKNVEGVYKCQCMEGFTFDKEKNVCLSSAGENLYFQGSSSHHHHHHHHGGGSAWSHPQFEK

Perpule:alpha-factor signal peptide
Red:30K signal peptide
Green:His-tag
Light blue:StrepTagII
Orange:TEV protease recognition sequence(Q↓G)
Underlined:Pvs25

The alpha-factor signal peptide cleavage of PpPvs25(N-tag) occurs at ~~KR↓EAEA~~, EAEA is signal peptidase recognition region. The cleavage was determined as previously (Ref.23).

The 30K signal peptide cleavage of BmPvs25(N-tag) occurs at ~~NA↓HH~~,while that of BmPvs25(C-tag) occurs at ~~DA↓VT~~predicted by SignalP 6.0 server at https://services.healthtech.dtu.dk/service.php?SignalP.

Theoretical molecular weights of matured BmPvs25(N-tag) and BmPvs25(C-tag) are 22662.43 Da and 22868.63 Da, respectively, which were calculated by Compute pI/Mw tool at https://web.expasy.org/compute_pi/.

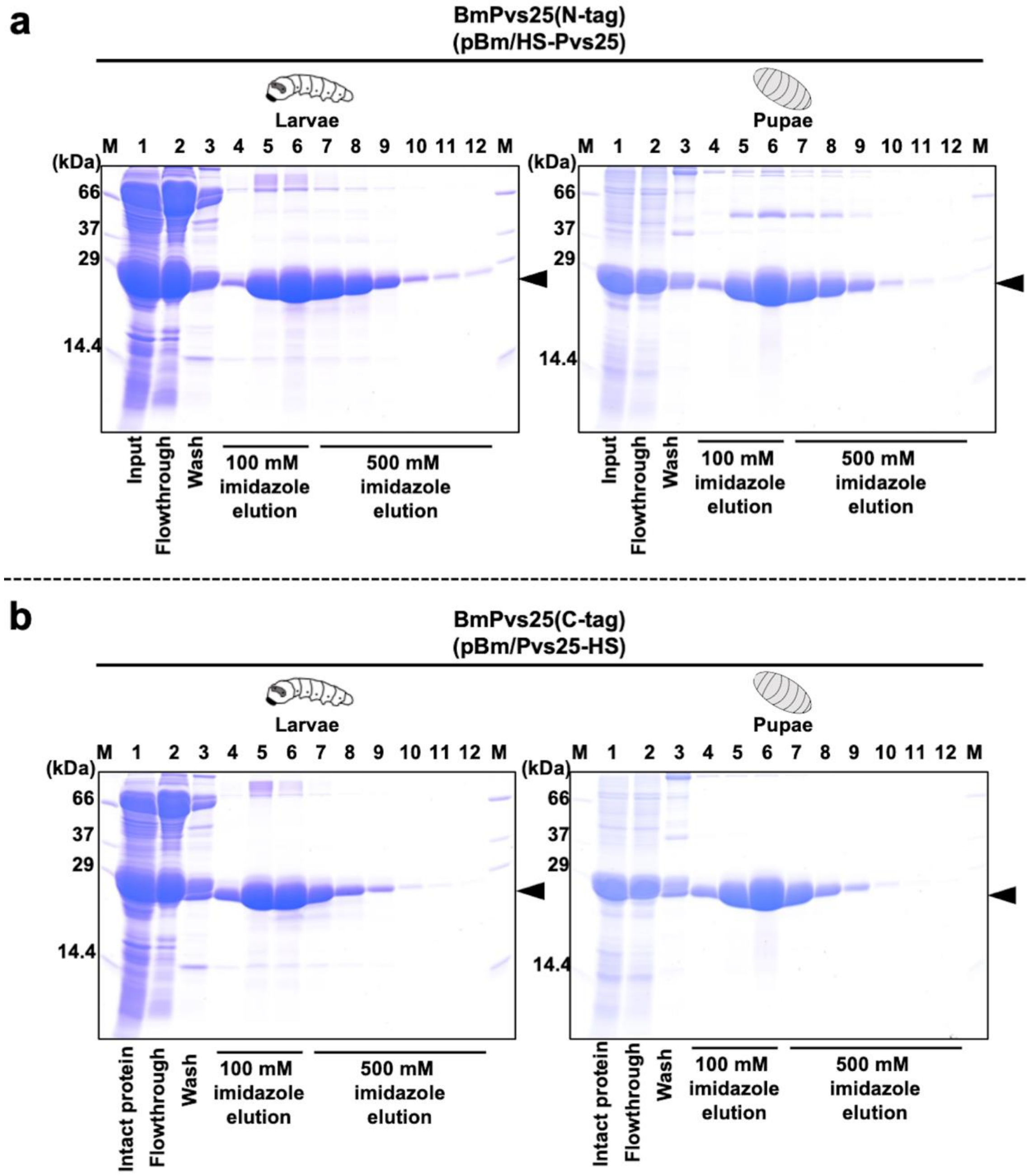


**Figure S1**. SDS-PAGE analysis on the fractions of Ni-affinity chromatography of BmPvs25s. a: Purification results of BmPvs25(N-tag) expressed in silkworm larvae and pupae. b: Purification results of BmPvs25(C-tag) expressed in silkworm larvae and pupae.

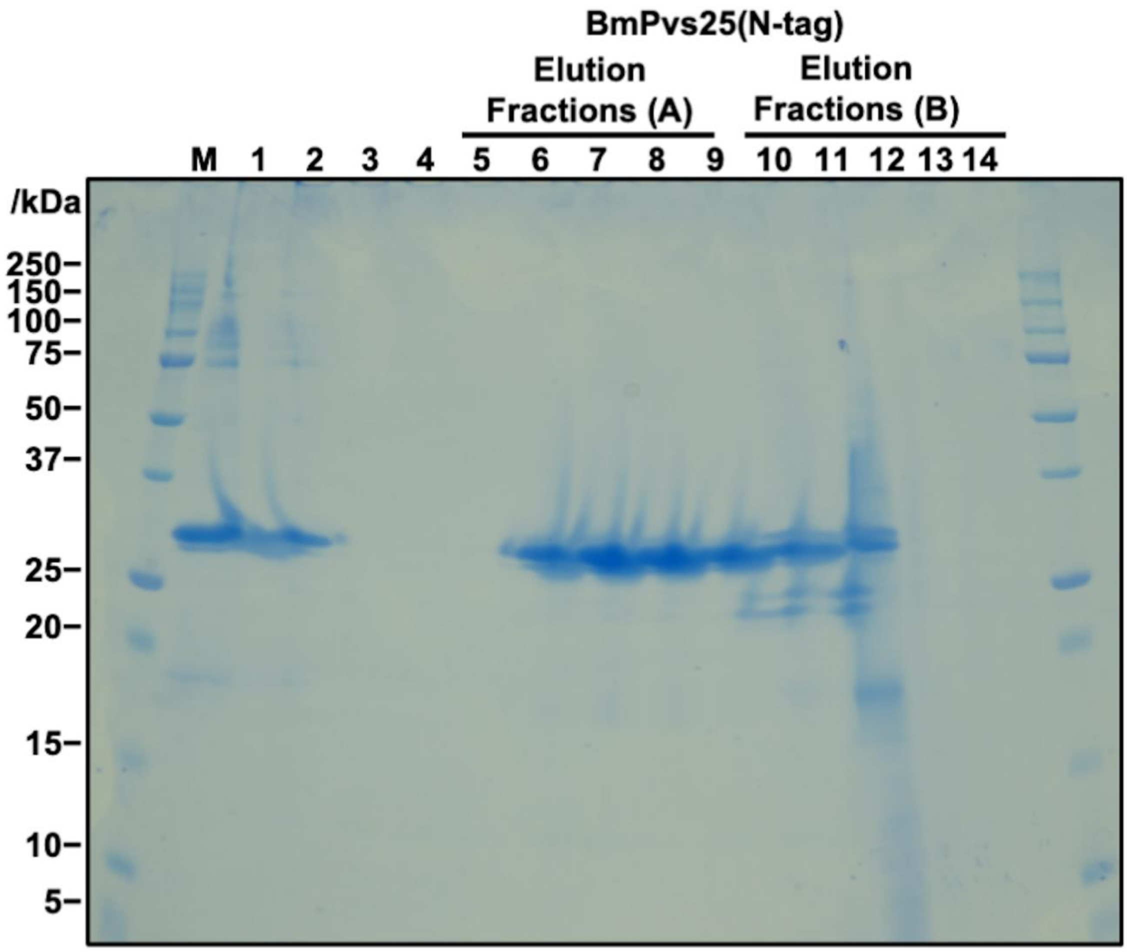


**Figure S2.** Hydrophobic interaction chromatography of BmPvs25(N-tag) from silkworm larvae Elution fractions (A) and elution fractions (B) are eluates of PBS. Each fraction was subjected to SDS-PAGE (15% acrylamide)/CBB stain. M: molecular mass, lane 1: His-tag purified protein, lane 2: His-tag purified protein diluted with $NH_4SO_4$ (final 2 M), lane 3: flowthrough fraction, lane 4: wash fraction, lanes 5-9: elution fraction with 2 M $NH_4SO_4$, lanes 10-14: elution fractions with PBS.

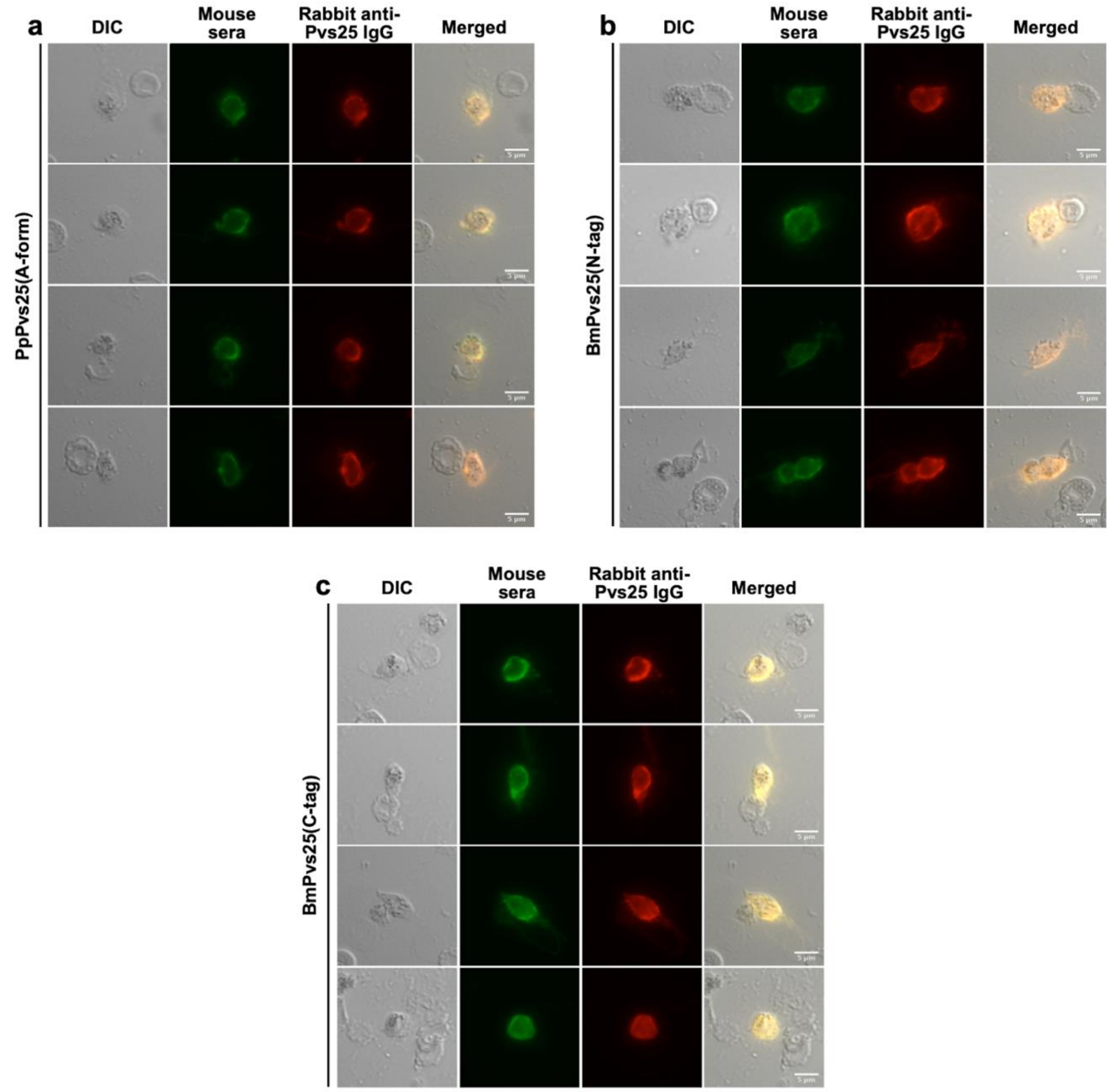


**Figure S3.** Additional images for the results shown in Figure 5 in the manuscript. a: immunostaining results using mice sera immunized with PpPvs25(A-form). b: immunostaining results using mice sera immunized with BmPvs25(N-tag). c: immunostaining results using mice sera immunized with BmPvs25(C-tag). Zygotes and ookinetes (retorts and mature ookinetes) cultured *in vitro* were fixed with acetone. The immunostaining results by using mouse sera induced by PpPvs25(A-form), BmPvs25(N-tag), or BmPvs25(C-tag) were indicated as green color (Alexa Fluor 488). Immunostaining results using rabbit antibody specific to Pvs25 were indicated as red color (Alexa Fluor 546). Merged, merged image of green and red signal. Bars, 5 µm.
